# Somatic gene repression ensures physical segregation of germline and soma in *Drosophila* embryos

**DOI:** 10.1101/2025.06.13.659480

**Authors:** Miho Asaoka, Mizuki Kayama, Tomoki Kawagoe, Makoto Hayashi, Shumpei Morita, Satoru Kobayashi

**Affiliations:** Life Science Center for Survival Dynamics, Tsukuba Advanced Research Alliance (TARA), University of Tsukuba, Tsukuba, Ibaraki, Japan; Graduate School of Life and Environmental Sciences, University of Tsukuba, Tsukuba, Ibaraki, Japan; Graduate School of Science and Technology, University of Tsukuba, Tsukuba, Ibaraki, Japan; Institute for Aquaculture Biotechnology (IAB), Tokyo University of Marine Science and Technology, Minato-ku, Tokyo, Japan; Asamushi Research Center for Marine Biology, Tohoku University, Aomori, Japan

**Keywords:** germ cell, cellular protrusion, germ/soma separation, cell death, Nanos, Pgc, germ plasm, *Drosophila*

## Abstract

In many animals, primordial germ cells are transiently segregated outside the somatic-cell cluster that forms the embryo’s body during early embryogenesis. This physical segregation of germline from the soma has been long believed to be crucial for germline development, but the mechanisms controlling this segregation and its developmental significance remain unclear. Here, in *Drosophila*, we show that somatic gene silencing in the germline is essential for maintaining this segregation. We showed that primordial germ cells (pole cells) lacking the Nanos- and Polar granule component (Pgc)-dependent dual repression mechanism caused misexpression of widespread somatic genes. They formed abnormal cellular protrusions, invaded adjacent somatic epithelium, and intermingled with somatic cells. These mislocalized pole cells ultimately underwent cell death, whereas properly segregated cells survived. Notably, the knockdown of *miranda* (*mira*), one of the somatic genes expressed ectopically in the pole cells lacking this repression mechanism, rescued these phenotypes. Our findings provide the first evidence that somatic gene silencing in germline is essential for preserving the physical segregation between germline and soma, highlighting its role in ensuring germline viability during early development.

## Introduction

The physical segregation of primordial germ cells from the soma is widely observed in many animal species [1]. For example, in some hymenopterans and higher dipterans, including *Drosophila*, primordial germ cells (pole cells) are formed at the blastoderm stage by inheriting a specific ooplasm called germ plasm, which contains maternal factors required for germline development. They are formed outside the somatic layer at this stage and remain outside the somatic cells until mid-embryogenesis. They reside inside the lumen of developing midgut primordium, which is derived from the outside of the somatic layer at the blastoderm stage. Pole cells then pass through the midgut epithelium into the hemocoel and migrate into the embryonic gonads, where they differentiate into oocytes or sperms [2–5]. A similar spatial segregation is observed in some vertebrate species [6, 7]. In some reptiles and primates, primordial germ cells first appear in the extra-embryonic region, separated from the somatic-cell cluster forming an embryo [7–9]. In chicks and mice, primordial germ cells are initially observed inside the somatic-cell cluster but then migrate into the extra-embryonic region and stay there until mid-embryogenesis [10–12]. From these observations, the physical segregation of germline from the soma has been recognized as a common feature across species and is thought to play an important role in germline development [1]. However, its underlying mechanisms and developmental significance remain poorly understood. This is due to the lack of experimental model that disrupts this segregation.

Besides the physical segregation, somatic gene silencing in germline has also been reported in many animal species. Expression of the genes required for somatic tissue development are specifically suppressed in primordial germ cells during early embryogenesis [13–21]. Recently, in *Drosophila*, two maternal proteins, Nanos and Polar granule component (Pgc), has been identified as factors required to repress somatic gene expression in pole cells [13, 16]. Nanos, a translational repressor, suppresses the translation of *imp-α2* mRNA, which encodes an adaptor protein for nuclear import receptors. This results in the inhibition of Imp-α2-dependent nuclear import of transcriptional activators in pole cells [16]. Pgc suppresses RNA polymerase II-dependent transcription by inhibiting the function of positive transcriptional elongation factor B (P-TEFb) [22]. Nanos-dependent inhibition of the nuclear import of transcriptional activators and Pgc-dependent global transcriptional silencing are both required to suppress the expression of somatic genes, such as *fushi tarazu* (*ftz*), *even-skipped* (*eve*), and *Sex-lethal* (*Sxl*) in pole cells. Thus, Nanos-and Pgc-dependent repression acts as a ‘double-lock’ mechanism to ensure tight inhibition of somatic gene expression in pole cells [16].

Here, we describe the critical role of Nanos/Pgc-dependent somatic gene repression in the spatial segregation of the germline from the soma. We also present data suggesting that the loss of this mechanism reduces the number of germline progenitors.

## Results

### Transcriptome of pole cells lacking Nanos/Pgc-dependent double-lock mechanism

To comprehensively identify genes suppressed by the Nanos/Pgc-dependent ‘double-lock’ mechanism in pole cells, we first compared the transcriptomes of pole cells lacking the double-lock mechanism with those of normal pole cells. A previous study has shown that the double-lock mechanism is disrupted by overexpression of the Imp-α2 protein in pole cells mutant for maternal *pgc* (*pgc^-^ impα2OE* pole cells) [16]. Using fluorescence-activated cell sorting (FACS), we isolated pole cells from *pgc^-^ impα2OE* and normal (*w*) embryos at stages 4–6 and processed these cells for RNA-sequencing (RNA-seq). We found that, compared to normal pole cells, 878 and 398 transcripts were upregulated and downregulated in *pgc^-^ impα2OE* pole cells, respectively [False discovery rate (FDR) < 0.01] (Fig 1A). *fushi-tarazu* (*ftz*) and *Sex-lethal* (*Sxl*), of which expression are reported to be suppressed by the double-lock mechanism in our previous paper [16], were observed in the “upregulated transcripts”. Gene ontology (GO) terms related to somatic development, such as “tissue development” and “generation of neurons” were enriched in the upregulated transcripts (genes) (Fig 1B and S1 Table). Furthermore, 389 transcripts were upregulated specifically in *pgc^-^ impα2OE* pole cells, but not in *pgc^-^* or *impα2OE* pole cells (Fig 1C and S2 Table), and the GO terms related to somatic development were also enriched in them (Fig 1D and S2 Table). The above results indicate that widespread misexpression of “somatic genes” occurs in *pgc^-^ impα2OE* pole cells.

**Fig 1.**
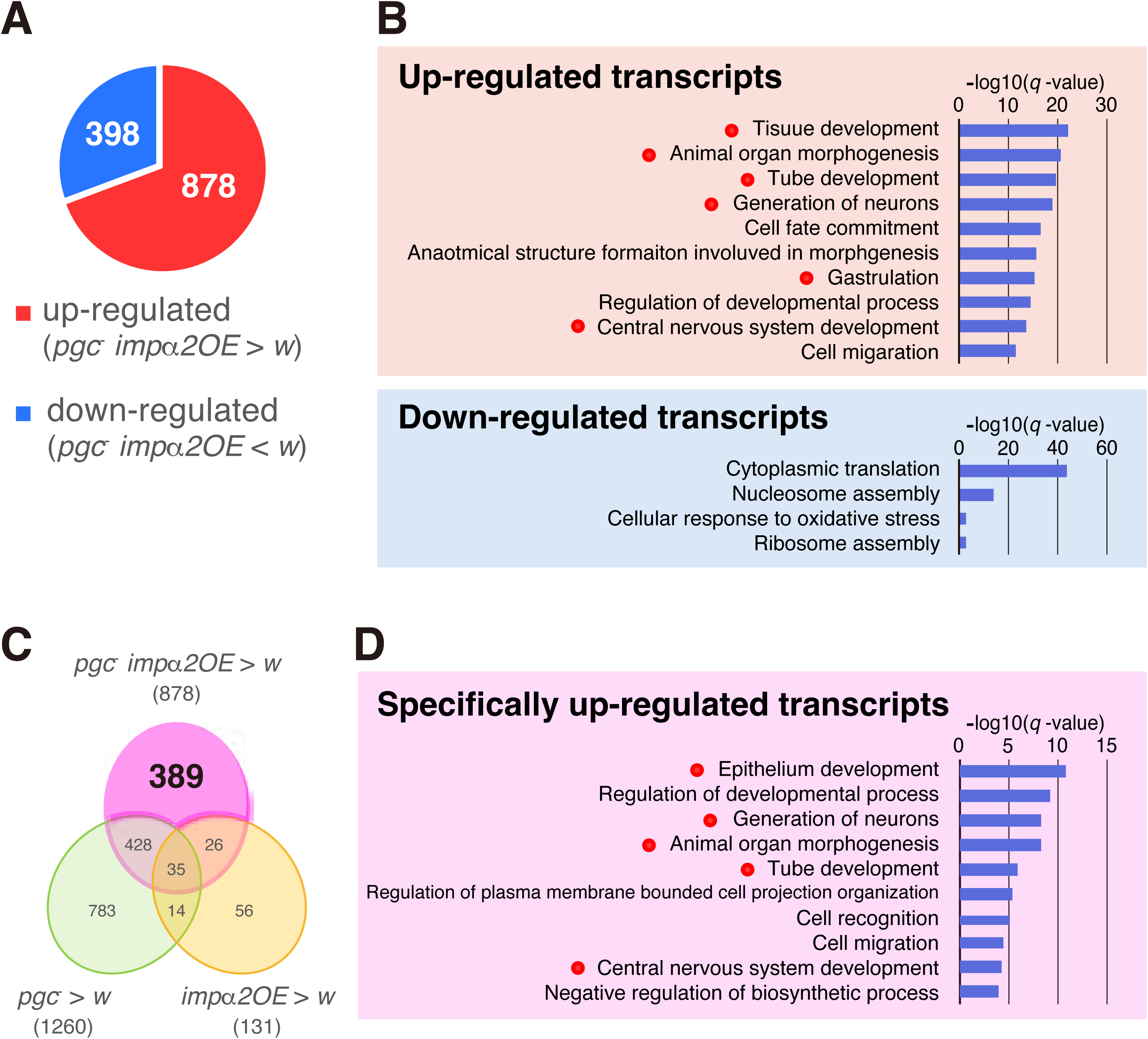
Transcriptomic profile of pole cells lacking the “double-lock mechanism.” (**A**) Number of transcripts upregulated and downregulated in *pgc^-^ impα2OE* pole cells compared with *w* pole cells (FDR < 0.01). (**B**) GO term enrichments of upregulated and downregulated transcripts shown in (**A**). Top 10 GO terms for upregulated transcripts are shown. All significantly enriched GO terms for downregulated transcripts are shown (*q* < 0.01). See also Tables S1 and S3. (**C**) Comparison of upregulated transcripts (FDR < 0.01, vs. *w*) between *pgc^-^*, *impα2OE*, and *pgc^-^ impα2OE* pole cells. The number of transcripts belonging to each category is shown. (**D**) GO term enrichment of 389 transcripts specifically upregulated in *pgc^-^ impα2OE* pole cells. See also Table S2. The GO terms related to the somatic tissue development are marked with red dots in (**B)** and (**D)**.

The transcripts downregulated in *pgc^-^ impα2OE* pole cells were enriched for GO terms unrelated to germline development (Fig 1B and S3 Table). Furthermore, we noticed that the expression of transcripts encoding germ-plasm components, such as *oskar* (*osk*)*, germ cell-less* (*gcl*)*, nanos, P-element induced wimpy testis* (*piwi*)*, Peroxiredoxin 2* (*Prx2*)*, Trapped in endoderm1* (*Tre1*), and *wunen2* (*wun2)*, was not downregulated in *pgc^-^ impα2OE* pole cells (S4 Table), suggesting that maternal transcripts encoding germ-plasm components were normally retained in *pgc^-^ impα2OE* pole cells.

### Morphological alterations of *pgc^-^ impα2OE* pole cells

In normal embryos, pole cells are formed outside the somatic-cell layer at the blastoderm stage (stage 4), and they remain in the lumen of the posterior midgut primordium until the end of gastrulation (stage 9) (Fig 2A). During these stages, the pole cells exhibit a spherical morphology [2] (Fig 2B). In contrast, *pgc^-^ impα2OE* pole cells formed cellular protrusions at the onset of gastrulation at stage 6. A single protrusion was formed per cell and penetrated between the epithelial cells (Fig 2B and S1A Fig). The percentage of *pgc^-^ impα2OE* pole cells forming protrusions was significantly higher than normal, *impα2OE,* or *pgc^-^* pole cells (Fig 2C). At stages 7–9, *pgc^-^ impα2OE* pole cells were located within the epithelial layer of the midgut primordium, whereas normal pole cells were in the lumen of the midgut primordium (Fig 2A, 2D). At these stages, *pgc^-^ impα2OE* pole cells were never observed in the hemocoel outside the epithelial layer of the midgut primordium (S1B Fig). The percentage of *pgc^-^ impα2OE* pole cells within the epithelial layer was significantly higher than that of normal, *impα2OE*, or *pgc^-^* pole cells (Fig 2E). Our observations strongly suggest that *pgc^-^ impα2OE* pole cells with protrusions invade the epithelial layer of the midgut rudiment and are intermingled with somatic cells.

**Fig 2.**
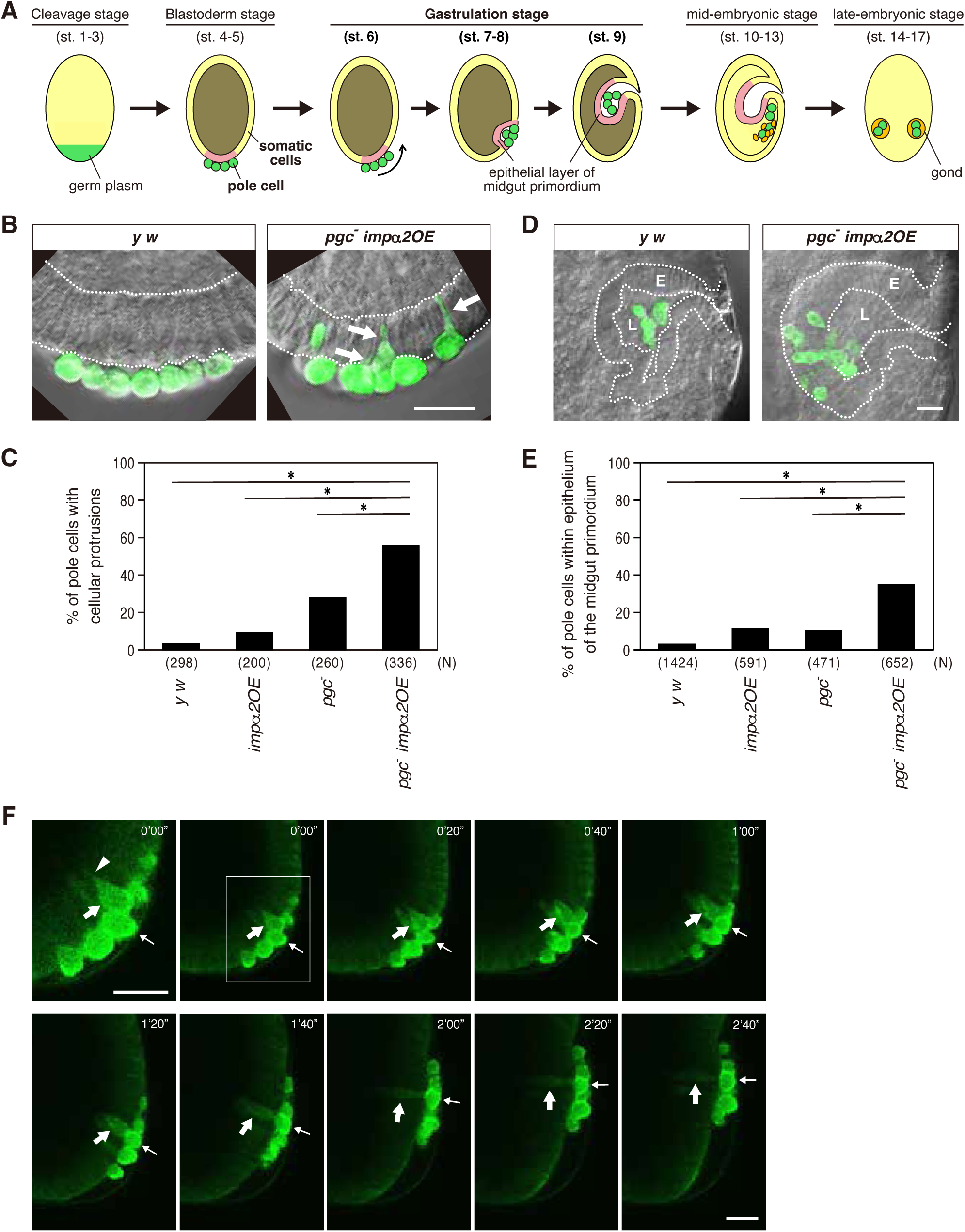
Pole cells lacking the double-lock mechanism form cellular protrusions and invade the somatic-cell layer. (**A**) Illustration of pole cell development in the *Drosophila* embryo. Germline, somatic cells, and yolk are shown in green, yellow, and brown, respectively. Somatic cells that give rise to the midgut and somatic gonadal precursors are shown in pink and orange, respectively. (**B**) Posterior pole regions of normal (*y w*) and *pgc^-^ impα2OE* embryos at stage 6. Embryos were stained for the germline marker Vasa (green). DIC images merged with the Vasa signal are shown. White dotted lines outline the somatic-cell layer. Arrows indicate cellular protrusions. See also Figure S1. (**C**) Percentage of pole cells with protrusions in *y w*, *impα2OE*, *pgc^-^*, and *pgc^-^ impα2OE* embryos at stage 6. (**D**) Magnified confocal images of midgut primordium of *y w* and *pgc^-^ impα2OE* embryos at stage 7. DIC images merged with the Vasa signal (green) are shown. The white dotted lines outline the epithelium of the midgut primordium. L: lumen of the midgut primordium; E: epithelium of the midgut primordium. (**E**) Percentage of pole cells in the epithelium of the midgut primordium in *y w*, *impα2OE*, *pgc^-^*, and *pgc^-^ impα2OE* embryos at stages 7-9. (**F**) Sequential time-lapse confocal images of the posterior pole region of a *pgc^-^ impα2OE* embryo from the end of stage 6 to the beginning of stage 7. Embryo with *nanos-moesin-GFP* transgene (green) visualizing cell membrane of germline cells [23]. The image in the upper left corner is a magnified image of the white box in the second panel from the left (time = 0’00”). Arrowhead indicates cellular protrusion. Thick and thin arrows indicate a pole cell that formed a protrusion at time 0’0” and that with a spherical shape, respectively. Asterisks in (**C**) and (**E**) indicate significance (*q* < 0.01, two-sided Fisher’s exact test). “N” indicates the number of pole cells examined. Panels (**A)**, (**B)**, (**D)**, and (**F)**: Embryos shown with anterior up, posterior down, ventral left, dorsal right. Scale bars, 20 µm.

This idea is supported by our live imaging. To visualize the cell membrane of pole cells in living embryos, we expressed the Moesin-GFP fusion protein only in pole cells [23]. *pgc^-^ impα2OE* pole cells forming protrusions at stage 6 invaded the epithelial layer of the midgut primordium during gastrulation, whereas *pgc^-^ impα2OE* pole cells with spherical morphology did not invade, as far as we observed (we examined six embryos for 20 min) (Fig 2F).

### Developmental fate of *pgc^-^ impα2OE* pole cells

In our live imaging, Moesin-GFP expression was maintained in spherical pole cells (thin arrows in Fig 2F), whereas it gradually disappeared in pole cells invading the midgut primordium (thick arrows in Fig 2F). Furthermore, in fixed samples, we often observed cell debris lacking DNA staining in the midgut primordium of *pgc^-^ impα2OE* embryos. These cell debris were positive for the germline marker Vasa, indicating that they were derived from pole cells (Fig 3A). These observations led us to hypothesize that *pgc^-^ impα2OE* pole cells invading the midgut primordium undergo cell death. To test this, we examined the expression of a cell death marker, the cleaved form of the *Drosophila* effector caspase, Dcp-1. In normal embryos, cleaved Dcp-1 was never detected in pole cells during gastrulation [the number of embryos examined (n) = 17]. In contrast, cleaved Dcp-1-positive pole cells were observed in 56.3% of *pgc^-^ impα2OE* embryos (*P* _vs. normal_ < 0.01, Fisher’s exact test, n = 16) (Fig 3B). The percentage of *pgc^-^ impα2OE* pole cells expressing cleaved Dcp-1 was significantly higher than normal pole cells (Fig 3C). The *pgc^-^ impα2OE* pole cells with cleaved Dcp-1 signal were observed in the epithelial layer of the midgut primordium rather than in the lumen (Fig 3D). The above observations strongly suggest that *pgc^-^ impα2OE* pole cells invading the epithelial layer of the midgut primordium undergo cell death.

**Fig 3.**
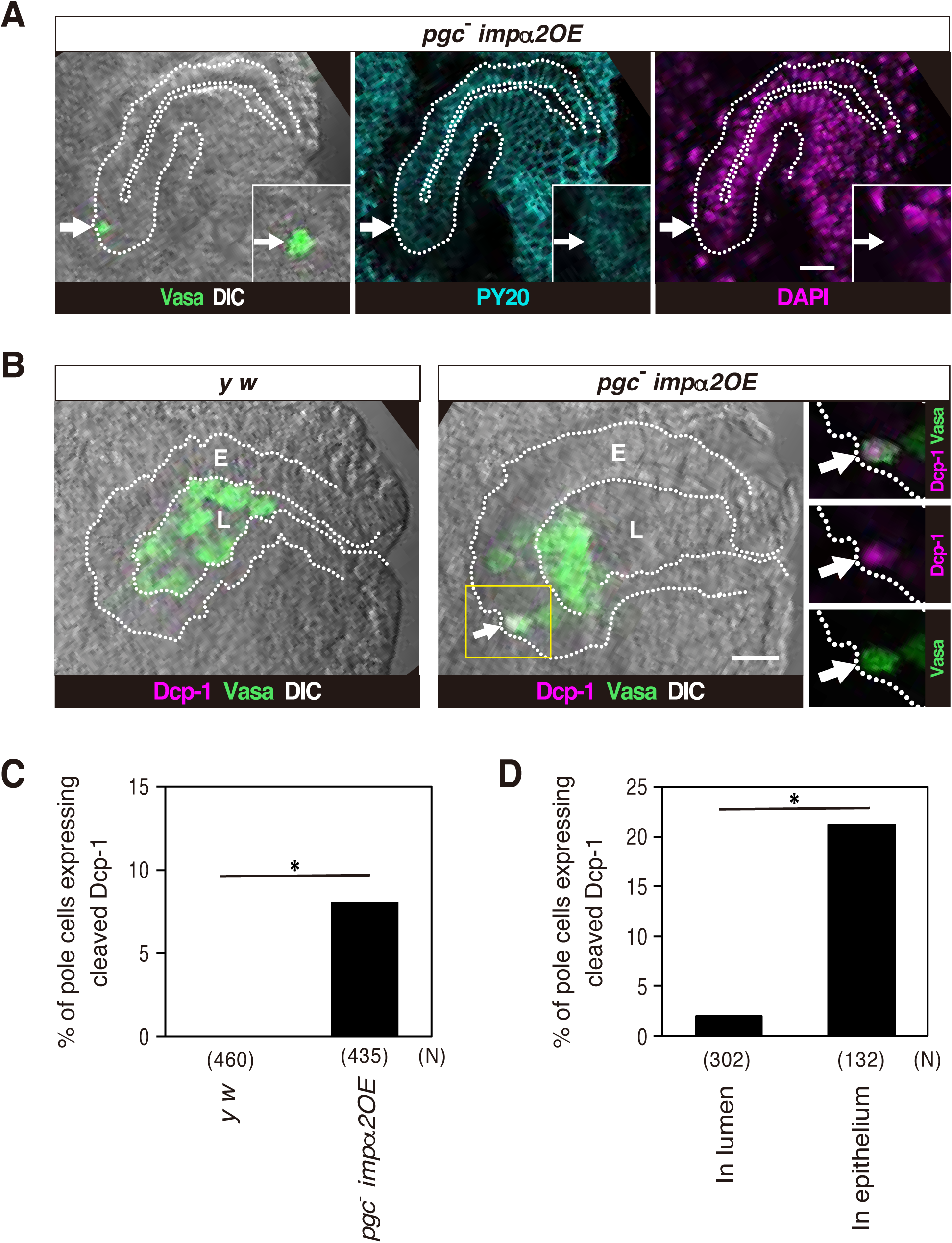
Pole cells lacking the double-lock mechanism express cleaved Dcp-1 in the midgut primordium epithelium. (**A**) Vasa-positive cell debris (arrows) in the *pgc^-^ impα2OE* embryo. Stage-9 embryo was triple-stained with anti-Vasa (green), and PY20 anti-phosphotyrosine (cyan) antibodies and DAPI (nuclear DNA, magenta). PY20 antibodies were used to outline all the cells [45]. DIC images merged with Vasa are also shown (left). Insets: Magnified images of the cell debris indicated by arrows. (**B**) Cleaved Dcp-1 expression in pole cells of *y w* and *pgc^-^ impα2OE* embryos at stage 7. DIC images merged with cleaved Dcp-1- (magenta) and Vasa-(green) signals. Arrows indicate a cleaved Dcp-1-positive pole cell. E: epithelium of the midgut primordium; L: lumen of the midgut primordium. Magnified images of the yellow square area are also shown on the right. (**C**) Percentage of pole cells expressing cleaved Dcp-1 at stages 7–9 in *y w* and *pgc^-^ impα2OE* embryos. (**D**) Percentage of pole cells expressing cleaved Dcp-1 at stages 7–9 in the lumen and epithelium of the midgut primordium of *pgc^-^ impα2OE* embryos. The white dotted lines in (**A)** and (**B)** outline the epithelium of the midgut primordia. Asterisks in (**C**) and (**D**) indicate significance (*P* < 0.01, two-sided Fisher’s exact test). “N” indicates the number of pole cells examined. Panels (**A**) and (**B**): Embryos shown with anterior up, posterior down; ventral left, dorsal right. Scale bars, 20 µm.

### Abnormalities in the morphology and developmental fate of *pgc^-^ impα2OE* pole cells were rescued by suppressing the somatic gene *miranda* (*mira*)

Next, we searched for the gene(s) responsible for the phenotypes of *pgc^-^ impα2OE* pole cells. We assumed that the misexpression of the somatic gene(s) was responsible for the phenotypes. Of the 878 transcripts upregulated in *pgc^-^ impα2OE* pole cells, we first selected 191 transcripts with the GO term “Tissue development” (GO 0009888) (Fig1B and S1 Table). Among them, we selected 27 transcripts with GO terms related to neuroblasts [“Generation of neurons” (GO 0048699) and “Central nervous system development” (GO 007417)] (S1 Table), because a similar cellular protrusion was reported in *Drosophila* quiescent neuroblasts [24]. Among them, the expression of transcripts from *mira*, *scute* (*sc*), and *tailless* (*tll*) was upregulated in *pgc^-^ impα2OE* pole cells at a higher level [log2 fold change > 2.0, average transcripts per million (TPM) > 40.0], and their expression levels were higher than those in *impα2OE* or *pgc^-^*pole cells (S1 Table). We finally focused on *mira* because the percentage of *pgc^-^ impα2OE* pole cells expressing *mira* was similar to that of pole cells forming protrusions (Fig 4A).

**Fig 4.**
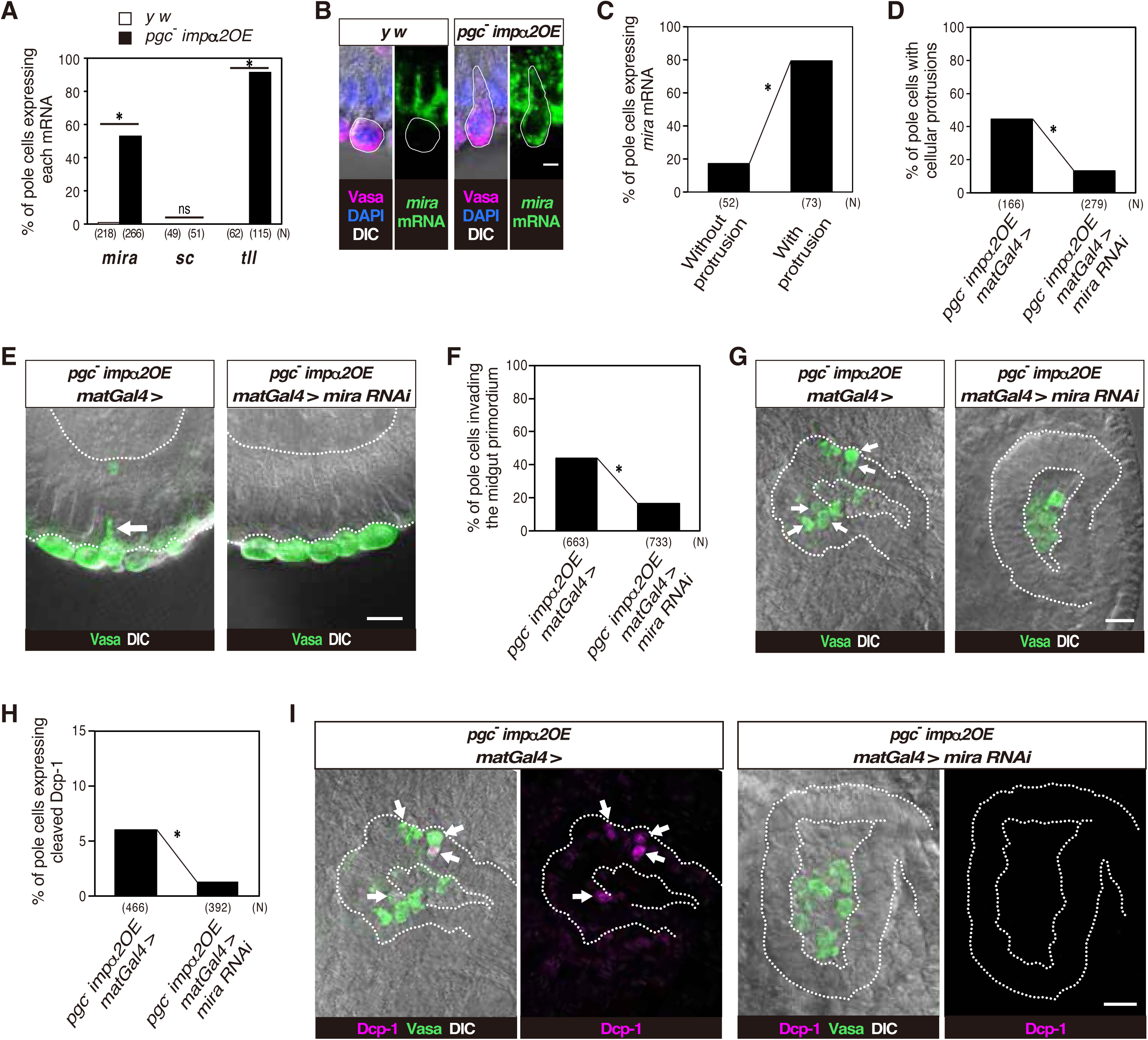
*mira* is one gene responsible for the phenotypes of *pgc^-^ impα2OE* embryos. (**A**) Percentage of pole cells expressing *mira*, *sc*, or *tll* mRNA in *y w* and *pgc^-^ impα2OE* embryos at stage 6. (**B**) *mira* mRNA expression in *y w* and *pgc^-^ impα2OE* embryos. Stage-6 embryos were triple-stained for *mira* mRNA (green), Vasa protein (magenta), and nuclear DNA (DAPI, blue). DIC images merged with Vasa and DAPI signals and images of *mira* mRNA signals are shown. White lines outline the pole cells. (**C**) Percentage of pole cells expressing *mira* mRNA in pole cells with or without protrusions in *pgc^-^ impα2OE* embryos at stage 6. (**D–I**) Phenotypes of protrusion formation, invasion, and cleaved Dcp-1 expression were examined in pole cells of *pgc^-^ impα2OE* embryos with (*pgc^-^ impα2OE matGal4 > mira RNAi*) or without knocking down *mira* (*pgc^-^ impα2OE matGal4 >*). Embryos were stained for Vasa (green) (**E** and **G**) or double-stained for Vasa (green) and cleaved Dcp-1 (magenta) (**I**). (**D**) Percentage of pole cells with protrusions at stage 6. (**E**) DIC images merged with the Vasa signal at stage 6. The arrow points to the pole cell with a protrusion. (**F**) Percentage of pole cells within the epithelium of the midgut primordium at stages 7–9. (**G**) DIC images merged with the Vasa signal (green) at stage 7. Arrows indicate pole cells invading the epithelium of the midgut primordium. (**H**) Percentage of pole cells expressing cleaved Dcp-1 at stages 7–9. (**I**) DIC images merged with cleaved Dcp-1- (magenta) and Vasa-(green) signals and Dcp-1-signal images are shown. Arrows indicate cleaved Dcp-1-positive cells. Asterisks in **(A**), (**C**), (**D**), (**F**), and (**H**) indicate significance (*P* < 0.01, two-sided Fisher’s exact test). “N” indicates the number of pole cells examined. The white dotted lines in (**E**) outline the somatic-cell layer, and those in (**G**) and (**I**) outline the epithelium of the midgut primordium. Scale bars: 5 µm (**B**), 10 µm (**E**), and 20 µm (**G**, **I**).

In normal embryos at stages 5–6, *mira* mRNA was detected in all somatic cells but not in pole cells (Fig 4A, 4B) [25–28]. Ectopic *mira* expression was observed in *pgc^-^ impα2OE* pole cells forming protrusions, but less frequently in spherical pole cells (Fig 4B, 4C). Thus, we wondered whether such ectopic *mira* expression in *pgc^-^ impα2OE* pole cells caused defects in morphology and developmental fate. We knocked down *mira* in *pgc^-^ impα2OE* pole cells by supplying double-stranded RNA (dsRNA) against *mira* using the *maternal-Gal4* (*matGal4*) driver. The frequency of *pgc^-^ impα2OE* pole cells with protrusions and those invading the epithelial layer of the midgut primordium was significantly reduced when *mira* dsRNA was expressed (Fig 4D-G). Furthermore, Dcp-1 expression in *pgc^-^ impα2OE* pole cells was reduced by *mira* dsRNA (Fig 4H, 4I). The above data showed that ectopic *mira* expression caused defects in *pgc^-^ impα2OE* pole cells.

### Ectopic *mira* expression was observed in a fraction of *pgc^-^ impα2OE* pole cells

In the above experiments, we noticed ectopic *mira* expression in a fraction of *pgc^-^ impα2OE* pole cells (Fig 4A). To examine the cause of the differences among these pole cells, we performed single-cell RNA-seq (scRNA-seq) using *pgc^-^ impα2OE* pole cells. Clustering analysis revealed that *pgc^-^ impα2OE* pole cells were divided into three clusters (clusters A, B, and C) (Fig 5A). Among them, *pgc^-^ impα2OE* pole cells in cluster C expressed *mira* at a higher level than those in clusters A or B (Fig 5B). In contrast, the expression of maternal transcripts localized in the germ plasm in cluster C was lower than that observed in clusters A or B (Fig 5C). These results suggest that *mira* is ectopically expressed in *pgc^-^ impα2OE* pole cells, which inherit fewer germ-plasm components than the other pole cells.

**Fig 5.**
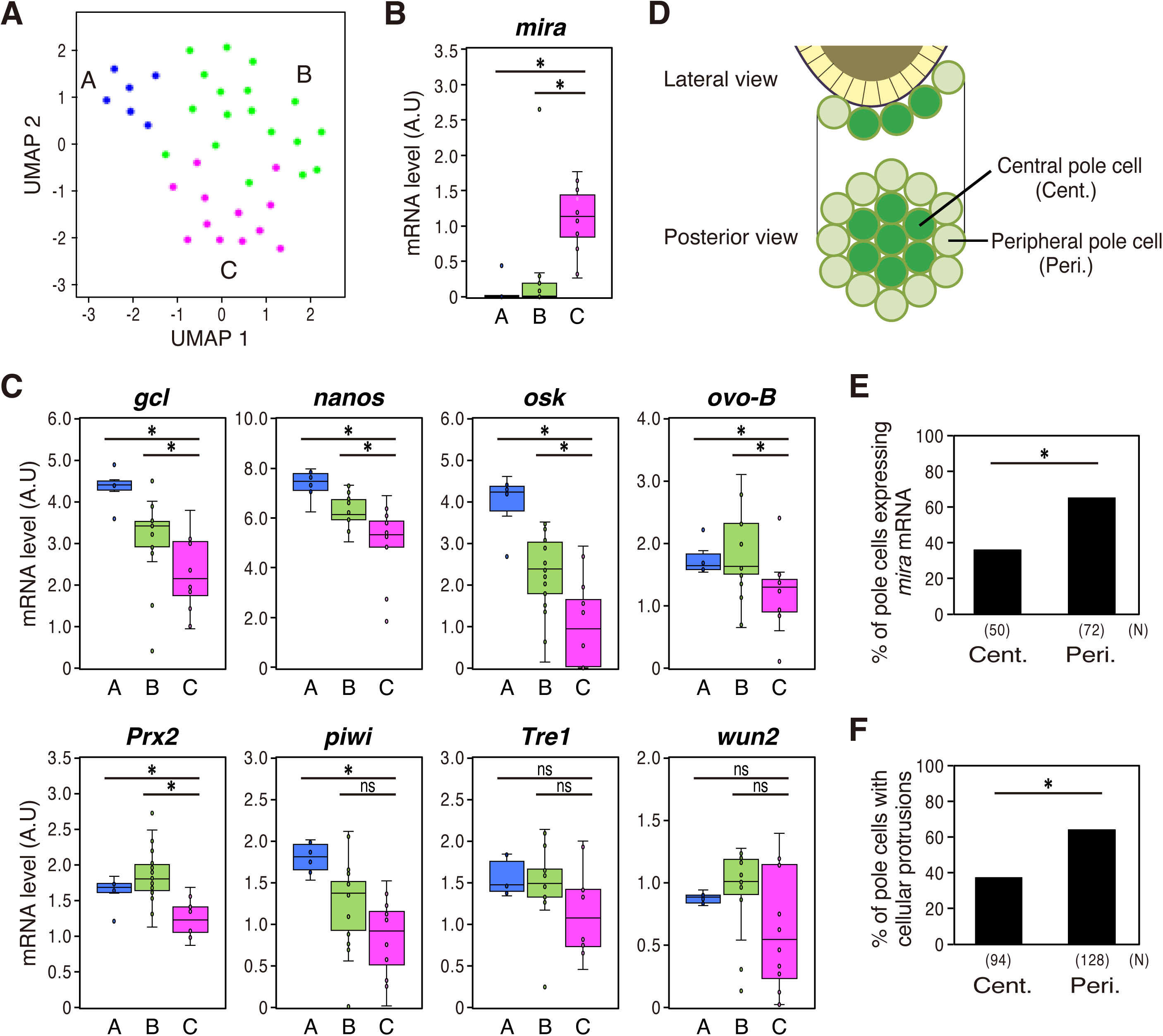
Heterogeneity in *pgc^-^ impα2OE* pole cells. (**A**) UMAP diagram showing three clusters from scRNA-seq of pole cells of *pgc^-^ impα2OE* embryos 2–3 h AEL (from the end of stage 4 to stage 6). (**B**-**C**) Box-and-whiskers diagram showing the distribution of expression levels of the indicated mRNA in pole cells belonging to clusters A, B, or C. The expression levels of *mira* mRNA (**B**) and eight major maternal mRNAs in the germ plasm (**C**) were examined. The “mRNA level” represents the normalized expression values (see Materials and Methods). See also Table S5. (**D**) Schematic drawing of central (green) and peripheral (light green) pole cells. Top: Lateral view of the posterior pole region of a stage-6 embryo. Bottom: a view from the posterior pole of the embryo. The ventral side was on the left and the dorsal side on the right. The pole cells were divided into two groups: peripheral (on the edge of the pole cell cluster) and central (the remaining cells). (**E**) Percentage of pole cells expressing *mira* mRNA in the central (Cent.) and peripheral (Peri.) pole cells of *pgc^-^ impα2OE* embryos at stage 6. (**F**) Percentage of pole cells with protrusions in the central and peripheral pole cells of *pgc^-^ impα2OE* embryos at stage 6. Significance was calculated using Steel’s method for multiple comparisons for cluster C in (**B)** and (**C)** [*: *P* < 0.05, ns (not significant): *P* > 0.05] and using the two-sided Fisher’s exact test in (**E)** and (**F)** (*: *P* < 0.05).

The pole cells located at the “central” region of pole cell cluster inherit a higher amount of germ plasm than those located in the “peripheral” region [29] (Fig 5D). Thus, we speculate that “peripheral” *pgc^-^ impα2OE* pole cells tend to misexpress *mira*. Figure 5E shows that the percentage of *pgc^-^ impα2OE* pole cells expressing *mira* was significantly higher in the “peripheral” than in “central” region. Similarly, the percentage of *pgc^-^ impα2OE* pole cells forming protrusions was also higher in the “peripheral” region (Fig 5F). Taken together, these results indicated that “peripheral” *pgc^-^ impα2OE* pole cells tended to misexpress *mira* and formed cellular protrusions.

## Discussion

In *Drosophila*, pole cells are never intermingled with somatic cells during cellularization, and this separation is maintained until mid-embryogenesis [2]. Here, we provide the first report of a molecular mechanism that controls the spatial separation between pole cells and the soma (Fig 6). Our results showed that *pgc^-^ impα2OE* pole cells formed cellular protrusions and invaded between epithelial cells of the midgut rudiment, resulting in the mingling of germline cells with the soma. We further showed that these phenotypes were rescued by knockdown of the somatic gene *mira*. Thus, repression of *mira* by Nanos-dependent inhibition of Imp-α2 production and Pgc-dependent suppression of RNA polymerase II activity is essential for the spatial segregation of the germline from the soma. In the absence of these two repression mechanisms, pole cells misexpressed *mira* and intermingled with the soma eventually die.

**Fig 6.**
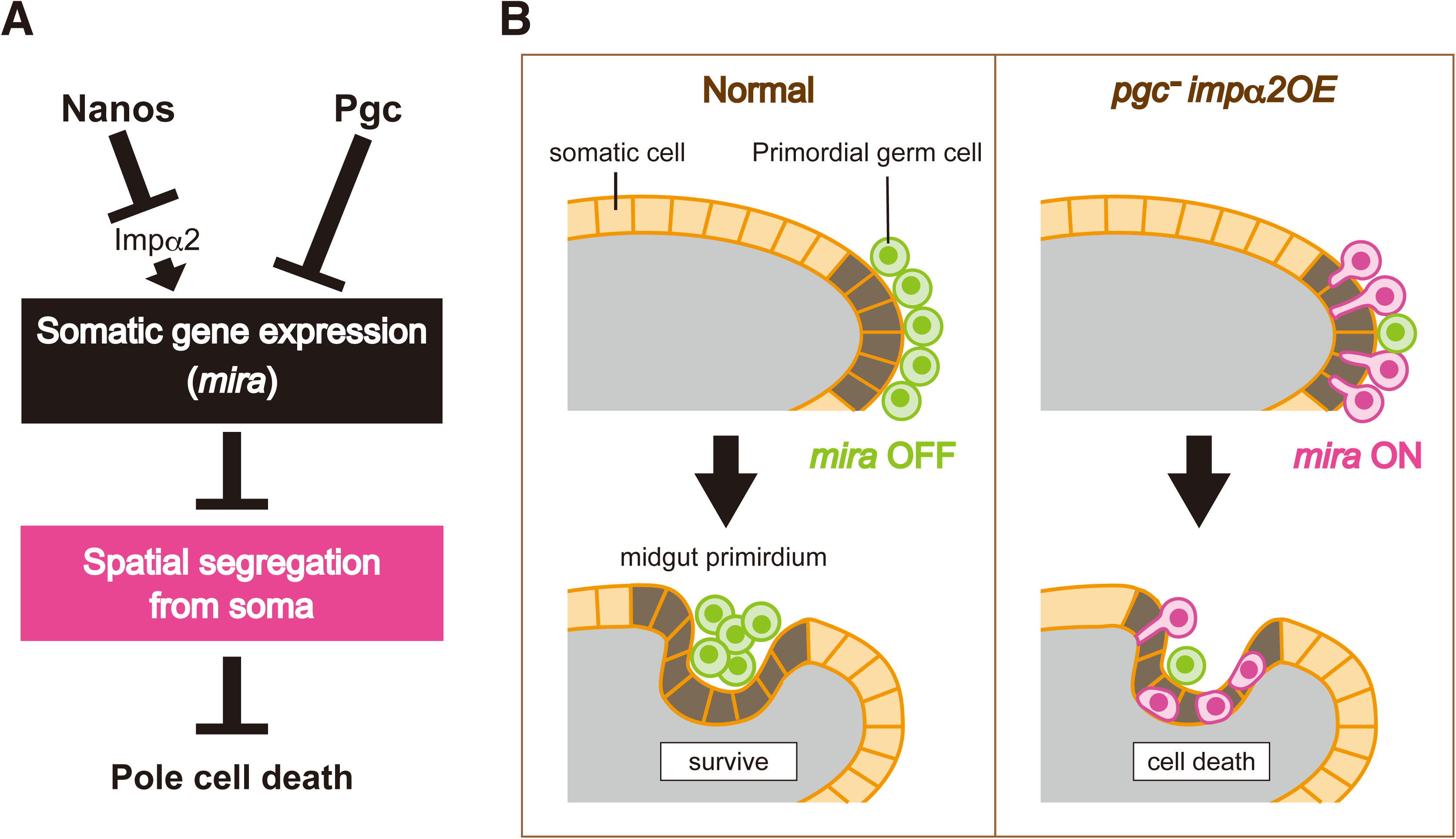
(**A**) Proposed model for the maintenance of germline/soma physical segregation. Nanos-dependent inhibition of Imp-α2 production and Pgc independently and redundantly repress expression of the somatic gene *mira* in pole cells. This suppression of *mira* permits pole cells remaining separated from the soma, and thereby preventing pole cell death. (**B**) In normal embryo (**Left**), pole cells do not express *mira*. They maintain their spatial segregation from the somatic layer during gastrulation. In contrast, in *pgc^-^ impα2OE* embryos (**Right**), *mira* (magenta) is misexpressed in a subset of pole cells. Mira-expressing pole cells form protrusions, invade into the somatic layer, and intermingle with the somatic cells. These pole cells eventually die in the somatic layer. Pole cells inheriting less germ plasm tend to misexpress *mira* and exhibit the phenotypes.

These two mechanisms act independently and redundantly to repress somatic gene expression [16]. Misexpression of *mira*, as well as protrusion formation and failure in germline/soma segregation, was enhanced in *pgc^-^ impα2OE* pole cells compared to *impα2OE* or *pgc^-^* pole cells (S1 Table). Perhaps *mira* is expressed in the germline later in normal development and that its premature expression causes the phenotypes observed in *pgc^-^ impα2OE* pole cells. However, *mira* transcript (s) and their protein product (s) are never detected in the germline throughout normal development [25, 30]. Thus, it is unlikely that premature expression of *mira* causes the phenotypes observed in *pgc^-^ impα2OE* pole cells. In somatic cells, *mira* transcript (s) and protein product (s) are expressed ubiquitously at the blastoderm stage, and their expression is gradually restricted to neuroblasts in the ectoderm and stem-like cells (interstitial cell precursors: ICPs) in the midgut primordium [30, 31]. These cells are reported to form protrusions, the morphology of which is similar to that of *pgc^-^ impα2OE* pole cells [24, 32]. Although it remains unknown whether protrusion formation requires *mira* expression in these somatic cells, it is possible that *mira* can induce cellular protrusions of pole cells. In neuroblasts, the Mira protein binds to the cargo to transport proteins asymmetrically within the cells [33, 34]. In pole cells, Mira misexpression may induce the asymmetric localization of protein (s) required for the formation of cellular protrusions.

Our results showed that *pgc^-^ impα2OE* pole cells detected in the epithelial layer of the midgut primordium expressed cleaved Dcp-1, which eventually led to cell death. It is unlikely that the misexpression of widespread somatic genes in the pole cells affects their cell identity and results in cell death. In *pgc^-^ impα2OE* pole cells, genes required for the development of different somatic tissues, such as neurons, gut, and tubular tissues, were simultaneously expressed within a single pole cell. However, this mixed expression was observed not only in the pole cells of cluster C that expresses *mira* and forms protrusions, but also in those of clusters A and B (S2 Fig). Therefore, it is reasonable to speculate that the cell death of *pgc^-^ impα2OE* pole cells is induced directly by *mira* misexpression, although no evidence suggests that this gene is required for cell death. Alternately, it is also possible that the cell death is caused by *mira* through mingling of pole cells with inappropriate cells, or somatic cells. For example, unfit somatic cells straying into ectopic locations are selectively eliminated by autophagy through interaction (s) with the surrounding cells [35]. In unfit *Drosophila* cells, autophagy upregulates the pro-apoptotic gene *head involution defective* (*hid*), which induces cleaved Dcp-1 expression [35]. The autophagy regulators *Atg1* and *hid* are activated in *pgc^-^ impα2OE* pole cells (Table S1), whereas their expression is undetectable in normal pole cells until gastrulation is complete [25, 36].

In *Drosophila*, the “central” population of pole cells inherits more germ plasm than the “peripheral” population at the cellular blastoderm stage [29]. Germ plasm contains maternal factors for pole cell formation and subsequent development [37–40]. Our results showed that misexpression of *mira*, as well as protrusion formation, was prominent in the “peripheral” region of *pgc^-^ impα2OE* pole cells. Based on this observation, we propose that Nos and Pgc are sufficient to repress *mira* in the “peripheral” region, while in the central region, maternal factor(s) for germline formation repress it, along with Nos and Pgc. Another idea to explain this phenomenon is that Nos and Pgc repress *mira* only in the “peripheral” pole cells, inheriting a small amount of germ plasm. In any case, this mechanism helps the peripheral pole cells to execute germline development. Consequently, this may increase progeny production.

The expression of Nanos homologues and global transcriptional silencing are widely observed in the early primordial germ cells of numerous animals [8, 41–44]. Furthermore, their segregation from the soma has also been reported in numerous species [1, 3, 6–8]. Further studies will provide new insights into the evolutionarily conserved roles of somatic gene repression in spatial segregation between the germline and soma.

## Data availability statement

Bulk and single cell RNA-seq data were deposited with links to BioProject accession No. PRJDB20673 and No. PRJDB20688 in the DDBJ BioProject database, respectively.

## Acknowledgements

We thank R. Lehmann, A. Nakamura, E. Gavis, B. M. Mechler, the Bloomington *Drosophila* Stock Center, and the Drosophila Genomics Resource Center for fly stocks and materials, and Editage (www.editage.jp) for English language editing. We also thank Y. Kozono for valuable discussion. This work was supported by Grant-in-Aid for Scientific Research from the Japan Society for the Promotion of Science (JSPS) (23K05778 to M.A., and 24H02030 to S.K.).

## Author contributions

Conceptualization, M.A.; Investigation, M.A., M.K., and T.K.; Methodology, M.A., T.K, S.M., and M.H.; Supervision, M.A., and S.K.; Writing – original draft, M.A.; Writing – review and editing, M.H., and S.K.

## Supplemental Information

**S1 Fig. Phenotypes of *pgc^-^ impα2OE* pole cells.** (**A**) Posterior pole region of a *pgc^-^ impα2OE* embryo double-stained with anti-Vasa antibody (green) and DAPI (magenta) at stage 6. A DIC image merged with the Vasa signal is also shown (left). Arrowheads indicate a pole cell with cellular protrusion. The main body of the pole cell with the nucleus (arrowheads) remained on the outside, but its protrusion penetrated the somatic-cell layer (arrows). The pole cell protrusion (arrows) never reached beyond the somatic-cell layer into the yolk. (**B**) Posterior-dorsal region of a gastrulating *pgc^-^ impα2OE* embryo (stage 6/7) triple-stained with anti-Vasa antibody (green), PY20 antibody (magenta), and DAPI (blue). A DIC image merged with the Vasa signal is also shown (left). Arrows indicate pole cells in the epithelium of the midgut primordium. Note they were intermingled with the epithelial cells in the midgut primordium. Bars, 10 µm.

**S2 Fig. Expression of somatic genes in individual *pgc^-^ impα2OE* pole cells.** Heatmap showing the expression levels of somatic genes in each pole cell of Clusters A, B, and C. Expression levels are shown for three *mira* transcripts (red), transcripts related to neural development (magenta), transcripts annotated with the GO term “gastrulation” (blue), “tube development” (green), and transcripts encoding germ plasm components (orange). Transcripts related to neural development are defined as those annotated with “generation of neurons” and/or “central nervous system development”. Only transcripts fulfilling the following criteria are shown: (1) listed as “up-regulated transcripts” in bulk RNA-seq analysis (S1 Table); (2) annotated with at least one of the four GO terms, “generation of neurons”, “central nervous system development”, “gastrulation” or “tube development”; (3) assigned exclusively to one of the three somatic GO categories (neural development, gastrulation, tube development); and (4) expression level >0.5 in at least one cell in this scRNA-seq dataset.

**S1 Table** Transcripts upregulated in *pgc^-^ impα2OE* pole cells compared with normal pole cells, related to Figures 1A and 1B.

**S2 Table** Transcripts specifically upregulated in *pgc^-^ impα2OE* pole cells, related to Figures 1C and 1D.

**S3 Table** Transcripts downregulated in *pgc^-^ impα2OE* pole cells compared with normal pole cells, related to Figures 1A and 1B.

**S4 Table** Expression of the maternal transcripts localized to germ plasm.

**S5 Table** Transcripts used for the calculation of the ‘mRNA level’ in Figures 5B and 5C.

## Methods

### Fly strains

Flies were maintained on a standard *Drosophila* medium at 25 °C. The following fly stocks were used: *y w*, *impα2-nos3’UTR* [16], *EGFP-vasa* [46], *pgc^Δ1^/CyO* [22], *Df(2R)X58-7/CyO* [Bloomington Drosophila Stock Center (BDSC) No. 283], *nanos-Moe.EGFP.nanos 3’UTR* (*nanos-moesin-GFP*) [23], *w*; P{matα4-GAL-VP16}V37* (*maternal-Gal4*, BDSC No. 7063), *y v; P{TRiP.JF01314}attP2/TM3* (*UAS-mira^RNAi^*, BDSC No. 31356). Embryos produced from *impα2-nos3’UTR/impα2-nos3’UTR*, *pgc^Δ1^*/*Df(2R)X58-7*, *pgc^Δ1^ impα2-nos3’UTR/pgc^Δ1^ impα2-nos3’UTR* females mated with *y w* were referred to as *impα2OE, pgc^-^, and pgc^-^ impα2OE* embryos, respectively.

### Immunostaining

Antibody staining of embryos was performed as previously described [16]. Briefly, embryos were dechorionated in a sodium hypochlorite solution and fixed in a 1:1 mixture of heptane and fixative [4% paraformaldehyde in PBS (130 mM NaCl, 7 mM Na_2_HPO_4_, and 3 mM NaH2PO4)] for 20 min. Vitelline membranes of the fixed embryos were removed by vigorous shaking in 1:1 heptane:methanol. The following antibodies were used: chick anti-Vasa antibody (1:2000, lab stock) [47], mouse anti-Phosphotyrosine (PY20) antibody (1:50, BD Transduction Laboratories, 610000), and rabbit anti-cleaved Drosophila Dcp1 (Asp215) antibody (1:50, Cell Signaling Technology, 9578). Signals were detected using Alexa Fluor 488–conjugated anti-chicken IgY antibody (1:400, Thermo Fisher Scientific, A-11039), Cy3-conjugated anti-mouse IgG antibody (1:500, Jackson ImmunoResearch Laboratory, 715-165-151), Cy3-conjugated anti-rabbit IgG antibody (1:500, Jackson ImmunoResearch Laboratory, 711-165-152), or Alexa Fluor 633-conjugated anti-mouse IgG antibody (1:400, Thermo Fisher Scientific, A-21050), as appropriate. Nuclei were visualized in embryos by staining with DAPI (2 μg/ml, Sigma-Aldrich) for 30 min after antibody staining. Stained embryos were washed three times with PBT and mounted in VECTASHIELD with DAPI Mounting Medium (Vector Laboratories, H-1200).

Z-stack confocal images were taken using a Leica TCS-SP8 confocal microscope (Leica) and analyzed using Fiji software. For analysis of cellular protrusion formation of pole cells, z-stack images were taken with the following settings: x 63 objective lens, x2.5 digital zoom, 1024 × 1024 dpi, 0.36 μm z-step interval. The acquired confocal serial images were reconstructed into 3D structures and analyzed using Imaris software (ver. 7.6) (Oxford Instruments). The number of pole cells with protrusions was counted using the Fiji software.

### *In situ* hybridization

cDNA fragments corresponding to *sc* and *tll* were amplified from an imaginal disc cDNA library and an embryonic cDNA library [48] using the following primers: 5′-TCA TCG AGT GTG CTG TCC AC -3′ and 5′-TAG CTG AAG TTG GGA GTG CG -3′ for *sc*, and 5′-GGA GAT CCC GGC AGT ATG TG -3′ and 5′-GAG GGA TGG GTC CTC TGG AT -3′ for *tll*.

Amplified cDNAs were cloned into pGEM-T Easy (Promega, A1360). Templates for RNA probes were amplified from these plasmids and a full-length *mira* cDNA clone (BDGP Gold cDNA Collection, LD02989) using T7 and SP6 primers, and digoxigenin (DIG)-labeled RNA probes were synthesized from the fragments using T7 and SP6 RNA polymerase (Merck, 10881767001 and 10810274001), respectively.

The whole mount *in situ* hybridization of embryos was performed as previously described [16]. Triple staining for *in situ* hybridization, immunostaining with anti-Vasa antibody, and DAPI staining, as shown in Figures 4B, 4C, and 5E, were performed as follows. Embryos were dechorionated, fixed, and devitellinized as described in the **Immunostaining** section. All the following steps were performed using RNase-free reagents until the hybridization step was completed. Devitellinized embryos were immediately rinsed with methanol twice, dehydrated by incubating in ethanol four times (10 min each), incubated in 1:1 xylene:ethanol for 30 min, and rinsed with ethanol five times and with methanol twice (30 s each). The embryos were re-fixed in a 1:1 mixture of methanol and fixative for 5 min and washed twice with PBTw (PBS containing 0.1% Tween 20). The embryos were digested with 4 μg/mL Proteinase K in PBTw for 7 min at 23 °C. The reaction was stopped by washing the samples four times (2 min each) with PBTw. The embryos were post-fixed in fixative for 20 min, washed with PBTw five times (2 min each), incubated in a 1:1 mixture of PBTw and hybridization solution [HS; 50% formamide, 5 × SSC (750 mM NaCl and 75 mM sodium-citrate), 50 μg/mL heparin, 100 μg/mL yeast tRNA, 10 mM DTT, and 0.1% Tween 20] for 10 min, incubated in HS for 10 min, and then pre-hybridized in HS for 1.5 h and hybridized in HS containing 2 ng/μL RNA probe for 16 h at 56 °C. After hybridization, the embryos were washed four times with the washing solution (50% formamide, 5 × SSC, and 0.1% Tween 20) for 30 min each at 56 °C, with PBTw containing 75%, 50%, and 25% washing solution, and five times with PBTw for 5 min each at room temperature. The embryos were then blocked for 1 h in PBTB (PBS containing 0.5% Blocking Reagent [PerkinElmer, NEL700001KT] and 0.1% Tween 20), incubated in PBTB containing chick anti-Vasa antibody (1:1000) and anti-Digoxigenin-POD antibody (1:500, Roche, 11633716001) overnight at 4 ℃, and washed with PBTw once for 30 s and five times for 10 min each. The anti-digoxigenin-POD signal was detected using the TSA Plus fluorescein kit (PerkinElmer Life Sciences, Inc., NEL741001KT), and the embryos were washed with PBT (PBS containing 0.2% BSA and 0.1% Triton X-100) three times (15 min each), blocked for 30 min in blocking solution (PBS containing 0.2% BSA, 0.1% Triton X-100, and 5% normal goat serum), and incubated with Cy3-conjugated anti-chicken IgY antibody (1:250, Jackson ImmunoResearch Laboratory, 703-165-155) and DAPI for 2 h. After washing with PBT twice (30 s each) and three times (10 min each), the samples were mounted in VECTASHIELD with DAPI Mounting Medium. Z-stack confocal images were taken using a Leica TCS-SP8 confocal microscope (Leica) and analyzed using Fiji software.

### Live imaging of embryos

Embryos produced from *w*; *pgc^Δ1^ impα2OE; nanos-moesin-GFP* females mated with *y w* were collected at 1.5–3.5 h after egg laying (AEL), dechorionated in a sodium hypochlorite solution, and washed several times with distilled water (DW). Using forceps, embryos were aligned in a drop of PBS on a glass-bottomed dish (MATSUNAMI, D11130H) and covered with a piece of thin agarose gel (1% agarose in PBS, 1 mm thick). Time-lapse images were acquired using a Leica TCS-SP8 confocal microscope (Leica).

### *mira* knockdown

Embryos produced from *pgc^Δ1^ impα2-nos3’UTR/pgc^Δ1^ impα2-nos3’UTR; maternal-Gal4/UAS-mira^RNAi^* and *pgc^Δ1^ impα2-nos3’UTR/pgc^Δ1^ impα2-nos3’UTR; maternal-Gal4/+* females mated with *y w* were used as *pgc^-^ impα2OE* embryos with and without *mira* knockdown, respectively.

### Bulk RNA-seq analysis

Embryos produced from *w*; +; *EGFP-vasa, w*; *pgc^Δ1^ impα2-nos3’UTR/impα2-nos3’UTR; EGFP-vasa/+*, and *w*; *pgc^Δ1^ impα2-nos3’UTR; EGFP-vasa/+* females mated with *y w* were used as normal (*w*), *impα2OE*, and *pgc^-^ impα2OE* embryos, respectively. One hundred pole cells were isolated from 2–3 h AEL embryos by FACS, and cDNAs were synthesized using the SMART-Seq v4 Ultra Low Input RNA Kit for Sequencing (Clontech, 634890), as described previously [49, 50]. Nextera XT library creation and RNA-seq were performed at the University of Minnesota Genomics Center using the HiSeq 2500 platform (Illumina), and approximately 20 million reads per sample (50-bp paired-end reads) were obtained. Three biological replicates were generated and sequenced for each genotype. Raw read count data deposited with links to BioProject accession No. PRJDB8823 in the DNA Data Bank of Japan (DDBJ) BioProject database [49], were used as RNA-seq data from *pgc^-^* pole cells and re-analyzed in the same way as for other genotypes. The data obtained from normal, *impα2OE*, and *pgc^-^ impα2OE* pole cells were deposited with links to BioProject accession No. PRJDB20673 in the DDBJ BioProject database.

Raw reads were processed using Trimmomatic 0.36 [51] and aligned with the transcript model of *Drosophila melanogaster* (Flybase; *dmel-all-transcriptr-r6.17. fasta*) using the Kallisto software (ver. 0.44.1) with default settings [52]. Differential expression analysis was performed using R (ver. 4.4.1) using the edgeR package (ver. 4.2.1) [53]. For quality control of our dataset, the transcripts showing CPM (count-per-million) > 0.5 in at least three samples were filtered and then normalized by the trimmed-mean method before differential expression analysis. Differentially expressed transcripts were identified by comparing the normal (*w*) and experimental groups, considering batch effects to ensure accurate identification. Transcripts with an FDR, Benjamini-Hochberg procedure) of < 0.01 were determined to be differentially expressed transcripts.

### Gene enrichment analysis

GO functional enrichment analysis was performed using Metascape (ver. 3.5) (http://metascape.org). Upregulated and downregulated transcripts were analyzed separately.

### ScRNA-seq analysis

Embryos produced from *w*; *pgc^Δ1^ impα2OE; EGFP-vasa/*+ females mated with *y w* were used. Pole cells were isolated from 2–3 h AEL embryos and collected individually into different tubes via FACS, as described [50]. cDNAs were synthesized using the SMART-Seq HT Kit (Clontech, 634437), and libraries were prepared with 5 ng of cDNA input using the TruePrep DNA Library Prep Kit V2 for Illumina (Vazyme, TD502). The samples were sequenced using the HiSeq 10 platform (Illumina), and approximately 5 million reads per sample (150-bp paired-end reads) were obtained. Both library preparation and sequencing were performed at the Azenta Life Sciences. The raw fastq files were deposited with links to BioProject accession No. PRJDB20688 in the DDBJ BioProject database.

Raw reads were processed and aligned to the transcript model of *Drosophila melanogaster*, as described in the **Bulk RNA-seq analysis** section. The ‘est_count’ output generated by Kallisto was used as the raw read count for further processing in R (ver. 4.4.1) using Seurat (ver. 5.1.0) packages. For quality control of our dataset, transcripts showing a read count of > 0 in at least one sample were filtered and used. The feature counts were log-normalized and scaled using the default options. To remove dead cells, cells in which over 7.5% of the reads were from mitochondrial genes were filtered out. The final dataset of 36 high-quality cells and 21811 features was used for downstream analysis. To assemble cells into transcriptomic clusters using meaningful features, we first performed principal component analysis (PCA) using 2000 highly variable transcripts as input. Based on the Elbow method, the first eight principal component (PC) vectors were selected as significant PCs and used as input for Uniform Manifold Approximation and Projection (UMAP) clustering, with a resolution parameter of 1.0. Heatmaps in S2 Fig were generated using the ComplexHeatmap package (ver. 2.20.0) in RStudio (ver. 2024.09.1+394), based on log-normalized and scaled counts obtained from Seurat.

### Calculation of expression level of mRNAs of *mira* and components of germ plasm

To calculate the total number of transcripts corresponding to *mira* and germ-plasm components, normalized and scaled count data were used. For each gene, the counts of the transcripts shown in Table S5 were summed and used as the ‘mRNA level’ in Figures 5B and 5C.

### Statistics and reproducibility

The statistical tests used to analyze the data from each experiment are indicated in the figure legends. The experiments for all figures and tables were repeated more than twice, except for the scRNA-seq analysis shown in Figure 5.

